# Defects in lamin A-Prohibitin crosstalk lead to ROS elevation and OxPhos imbalance in laminocardiomyopathy

**DOI:** 10.1101/2025.07.17.665267

**Authors:** Subhradip Nath, Soumen Kanti Manna, Debasish Prusty, Sk. Ramiz Islam, Kaushik Sengupta

## Abstract

Lamins are critical for maintaining nuclear homeostasis, chromosome positioning, and cellular mechanotransduction, which involves the transfer of mechanical signals from the cellular microenvironment to the nucleus. Recent studies have also highlighted the involvement of lamin A in mitochondrial homeostasis and the regulation of reactive oxygen species production. Missense mutations in lamin A are linked to a spectrum of diseases known as laminopathies, which include conditions such as dilated cardiomyopathy (DCM), muscular dystrophy, and progeria. One such mutation, K97E, is associated with DCM, causing severe cardiac complications that can lead to myocardial infarction in extreme cases. Our study reveals a detailed pathogenic cascade in K97E-transfected cells involving disrupted interaction with Prohibitin-2, a key mitochondrial protein. Mitochondria exhibit increased fission, reduced fusion, and fragmentation, due to OPA1 downregulation and DRP1 recruitment driven by actin cytoskeletal remodelling. Impaired Rho-ERK– FAK signalling reduces F-actin assembly, elevating G-actin, which further promotes mitochondrial fission. This feedback loop leads to mitochondrial depolarisation, ATP deficiency, and global metabolic catastrophe, in particular cholesterol metabolism, accompanied by elevated ROS. In cardiomyocytes, such dysfunction may underlie contractile deficits and arrhythmias. Our findings establish PHB2 as a critical node linking nuclear integrity, cytoskeletal architecture, and mitochondrial homeostasis, offering new insights into DCM pathogenesis and therapeutic targets. Our findings elucidate the pivotal role of lamin A in cellular energetics and mechanotransduction, offering novel insights into DCM pathophysiology, which in turn opens avenues for developing targeted therapeutic strategies.

**Teaser:** Lamin A K97E mutation alters cellular metabolome through disturbed mitochondrial and actin homeostasis in a feedback loop with PHB2 at its hub and causes gross pathogenesis of DCM.

**Graphical Abstract:** Figure 5
Gross mitochondrial defects arising from PHB2 and actin perturbations leading to severe metabolic and bioenergetic effects during K97E mutation of lamin A

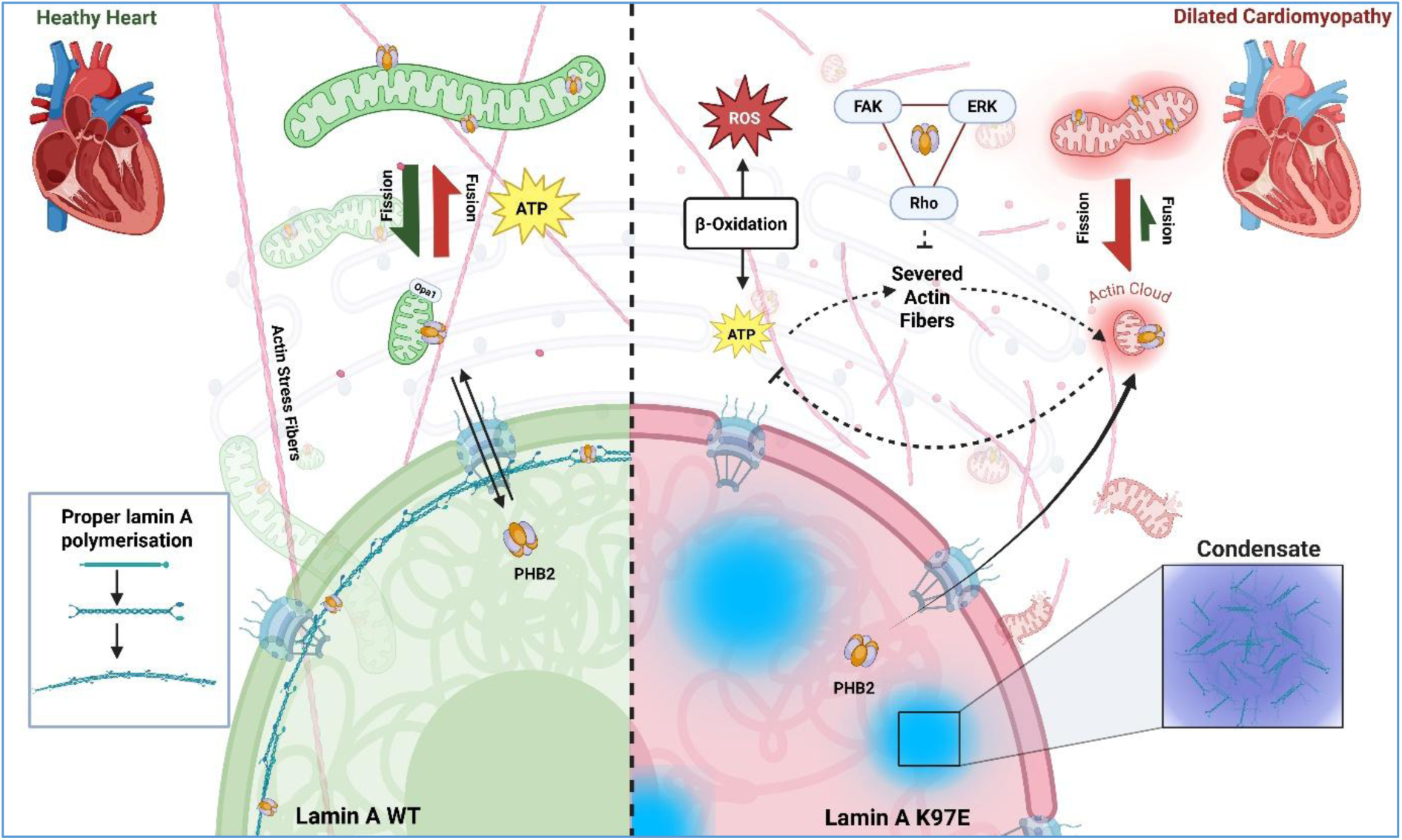

## Introduction

Lamins are type V intermediate filament proteins that form the fundamental structural component of the 14 nm lamina underlying the inner nuclear membrane. It plays a major role in maintaining proper nuclear architecture, genome organisation and epigenetic regulations (*1*). Encoded by the LMNA gene (1q22 in humans), lamin A and its splice variant lamin C contribute towards the dynamic structural framework of the nucleus, ensuring that nuclear shape and rigidity are preserved under various mechanical stresses. Recent cryo-electron tomographs have posited the presence of 3.5 nm lamin filaments, thereby shifting the paradigm of the formation of 10 nm fibres (*2, 3*). Lamin A is not only implicated in the maintenance of nuclear architecture and mechanical properties but also modulates cellular responses to the extracellular matrix, with its relative abundance varying according to tissue type and matrix stiffness through feedback from mechanotransduction signalling (*4, 5*). More than 500 LMNA mutations have been linked to 14 different human diseases, collectively known as ‘laminopathies’, including diseases like dilated cardiomyopathy (DCM), Emery-Dreifuss muscular dystrophy (EDMD), Hutchinson-Gilford progeria syndrome (HGPS), metabolic associated fatty liver disease (MAFLD), familial partial lipodystrophy type 2 (FPLD), and mandibuloacral dysplasia (MAD) (*1*). Laminopathic cells are characterised by an alteration of nuclear stiffness, fragile/misshapen nuclei, as well as altered chromosomal organisation as visualised by 3D-FISH experiments (*6*–*8*). In addition, epigenetic alteration has also been reported in several studies as one of the primary underlying causes of laminocardiomyopathy disorders (*9*–*11*). Among the laminopathies, DCM is the most prevalent laminopathic cardiac condition reported across the world, with 200+ associated LMNA mutations identified to date (http://www.umd.be/LMNA/). Lamin A-mediated DCM accounts for the pathogenesis of around 8-10% of overall DCM patients(*12*–*14*), primarily characterised by left ventricular dilation with impaired systolic functioning of the heart, resulting in sudden cardiac arrest and death in patients (*15*). Although the precise cellular mechanisms underlying DCM remain largely unknown, it is characterised by a gamut of pathophysiological alterations which include mitochondrial dysfunction, oxidative stress, cardiomyocyte apoptosis/necrosis, myocardial fibrosis, as well as lipotoxicity (*16*). Among these, mitochondrial dysfunction emerges as a critical factor, particularly in the context of heart failure, where an inadequate supply of ATP affects the contractile function of the heart. Mitochondria serve as a central hub for cellular metabolism, integrating multiple metabolic pathways to supply essential building blocks for macromolecule synthesis and the maintenance of metabolic homeostasis. Specifically, in cardiac tissue, the generation of ATP primarily depends on the fatty acid β-oxidation and the activity of the tricarboxylic acid (TCA) cycle: processes that are indispensable for sustaining the high energy demands of continuous cardiac contraction (*17*–*19*). Mitochondrial homeostasis is crucial for these processes and is maintained by dynamic processes including fission, fusion, mitophagy, and intracellular transport, which regulate mitochondrial morphology, integrity, abundance, and spatial distribution. In the event of fission, the outer mitochondrial membrane (OMM) constricts due to actin polymerisation within the endoplasmic reticulum (ER) contact sites, recruiting Dynamin-related protein 1 (Drp1) through adapter proteins such as mitochondrial fission 1 (Fis1) and mitochondrial fission factor (Mff)(*20*). This process segregates damaged components, allowing their selective removal via mitophagy, thereby preventing further cellular damage. On the other hand, mitochondrial fusion is a multistep process initiated by the activation of dynein-associated GTPases, including mitofusin (Mfn) 1/2 on the OMM and optic atrophy protein 1 (Opa1) on the inner mitochondrial membrane (IMM) (*21*). Following GTP hydrolysis-driven OMM and subsequent IMM fusion, mitochondrial contents are intermixed, diluting dysfunctional proteins and mtDNA to maintain homogeneity and functional stability (*22*). These processes are tightly regulated by various pathways and are activated according to cellular proliferation, differentiation and stress. Among their molecular gatekeepers, prohibitins (PHBs) play a pivotal role in maintaining fission-fusion balance and are also involved in mtDNA maintenance, oxidative phosphorylation or OxPhos assembly, cristae organisation, and apoptosis regulation (*23*–*25*). By stabilising Opa1, this complex promotes mitochondrial fusion, while its depletion results in mitochondrial fragmentation through improper Opa1 processing (*26*). Additionally, proper assembly of the PHB complex alongside m-AAA protease, Oma1, and cardiolipin maturation is critical for preserving mitochondrial dynamics, which can be hampered by alterations in the levels of either PHB1 or PHB2 (*27, 28*). Interestingly, PHB2 is also transported to the nucleus, where it interacts with various transcription regulators and chromatin remodelling proteins to modulate transcriptional activity, apoptosis, growth and differentiation in a tissue-specific manner (*29*–*31*). Recent studies have demonstrated that depletion of lamin A as well as its mutations results in increased reactive oxygen species (ROS) production and disrupts mitochondrial metabolic processes through multiple intermediary mechanisms (*32*–*35*). These findings highlight the importance of nuclear lamins in regulating mitochondrial function and conjecture potential novel avenues for delineating the link between metabolic abnormalities and the pathogenesis of DCM.

In this study, we focused on the K97E mutation in lamin A, which has been implicated in severe forms of dilated cardiomyopathy, particularly among several Italian families where affected individuals exhibited pronounced clinical symptoms (36). This mutation has been shown to disrupt the cellular epigenetic landscape and induce widespread changes in gene expression. In cultured cells expressing K97E lamin A, characteristic nuclear abnormalities and cytoplasmic aggregates of varying sizes and distributions are routinely observed (1, 37–39), likely resulting from enhanced lamin A self-association, as evidenced by calorimetric titration studies (37, 40). Our investigation revealed a profound reorganisation of the lamin A interactome, prominently involving Prohibitin 2 (PHB2), a key mitochondrial scaffold protein. This interactional shift contributes to mitochondrial dysfunction, manifesting as a gross collapse of cellular metabolism and bioenergetics. Specifically, we observed significant reductions in intracellular ATP levels, loss of mitochondrial membrane potential (depolarisation), and elevated production of reactive oxygen species (ROS), indicating severe mitochondrial impairment. We validated the nuclear interaction between lamin A and PHB2 and established its pathogenic relevance in the context of mitochondrial dysfunction in laminocardiomyopathy. In parallel, we identified marked disruptions in actin cytoskeletal organisation, characterised by diminished actin polymerisation and altered stress fiber morphology, driven by dysregulation of the ERK/Rho/FAK signalling axis. These cytoskeletal defects further contribute to mitochondrial dysfunction and reinforce the pathogenic cascade. Collectively, our findings demonstrate that the K97E lamin A mutation leads to interconnected mitochondrial and cytoskeletal abnormalities, establishing a mechanistic link between bioenergetic collapse and impaired mechanotransduction in the progression of DCM.

## Results

### Identification of PHB2-lamin A interaction and its disruption in the K97E mutation

K97E maps in the rod 1B domain (Fig. 1 A) of lamin A and is one of the most deleterious mutations of *LMNA,* causing DCM (*36*) with conduction defects. In retrospect, this mutant exhibited an increased propensity for self-association and a gradual disappearance from the lamina (*37, 38*). On logical grounds, we first questioned whether this enhanced self-association impacts the lamin A interactome. To address it, we employed proximity-based BioID2 labelling followed by mass spectrometric (MS) analyses (Fig. 1 B), where BioID2 is capable of biotinylating interacting proteins within a diameter of 20 nm. It is important to note that BioID has been acknowledged to be one of the key tools for studying the interactome of wild-type lamin A, as well as complex multimeric protein structures such as NPC and has promising applications in studying protein interactions in crowded environments (*39, 40*). The wild-type and mutant K97E-BioID2 plasmids were transfected into HEK293FT cell line, expressing a minimal amount of endogenous lamin A proteins and the cellular localisation was confirmed through immunofluorescence (Fig. 1 C, D). Along with the event of colocalisation of lamin A and myc-BioID2-lamin A variants (Fig. 1 E), it was confirmed that the tag was not affecting the physiological properties of lamin A, which showed similar distribution as reported in our earlier studies (*1*). An equal amount of lysate was used for streptavidin pull-down, followed by MS identification of the peptides (Fig. 1 F), which revealed a gross reduction in the number of interacting partners in case of K97E by half of the value compared to the wild type (Fig. 1 G). Among these, the mutant protein K97E exhibited a gain in interaction concerning 19 proteins and a loss of 100 proteins (Fig. 1 H). Upon enriching the set of proteins losing their interaction with mutant lamin A, we uncovered that the most prominent one was the PHB2 complex (Fig. 1 I), which is crucial for the maintenance of mitochondrial homeostasis, transportation, as well as regulation of growth, differentiation and apoptosis (*23, 25, 29, 41*). Further, we confirmed the loss of interaction of PHB2 with lamin A through BioID2-immunoblot (Fig. 1 J) as well as Co-IP (Supplementary Fig. 1). We chose C2C12 cells for all transfection-based studies as it has remained a faithful and established model system for studying cardiomyopathies (*42*). From immunoblot analysis, we observed a downregulation of PHB2 but not PHB1 in cells transfected by the mutant in both mRNA (Fig. 1 K) and protein levels (Fig. 1 L, M). Confocal images also revealed the same for PHB2, especially in terms of nuclear localisation (Fig. 1 N, O). To further validate these observations, we performed nuclear fractionation and found that indeed PHB2 levels were reduced to minuscule amounts in the nuclear extract of K97E-transfected cells (Fig. 1 P), pointing to a plausible perturbation of the interaction between K97E and PHB2. Theoretically, such a reduction in the amount of PHB2 in the nucleus should be compensated by an increase in the cytoplasmic fraction. To this end, the cytoplasmic extract of K97E-transfected cells was checked for PHB2 expression, which showed a modest increase (Fig. 1 Q). We asked ourselves whether it is a perturbation in the interaction between the mutant lamin A protein with PHB2, which in turn could retain PHB2 in the nucleus with much lower efficiency compared to the wild-type lamin A scenario. To check this hypothesis, we performed molecular docking studies using HADDOCK. For this, we utilized homology modelled full length PHB2 and lamin A proteins based on multiple PDB ids including 8J4I and 6JLB, for PHB2 (domain 1-190) and lamin A (domain 1-300) respectively (Supplementary Fig. 2). The energy parameters are shown in Table 1 which points to a distinctly weaker interaction of K97E with PHB2 (Fig. 1 R).

**Figure 1.**
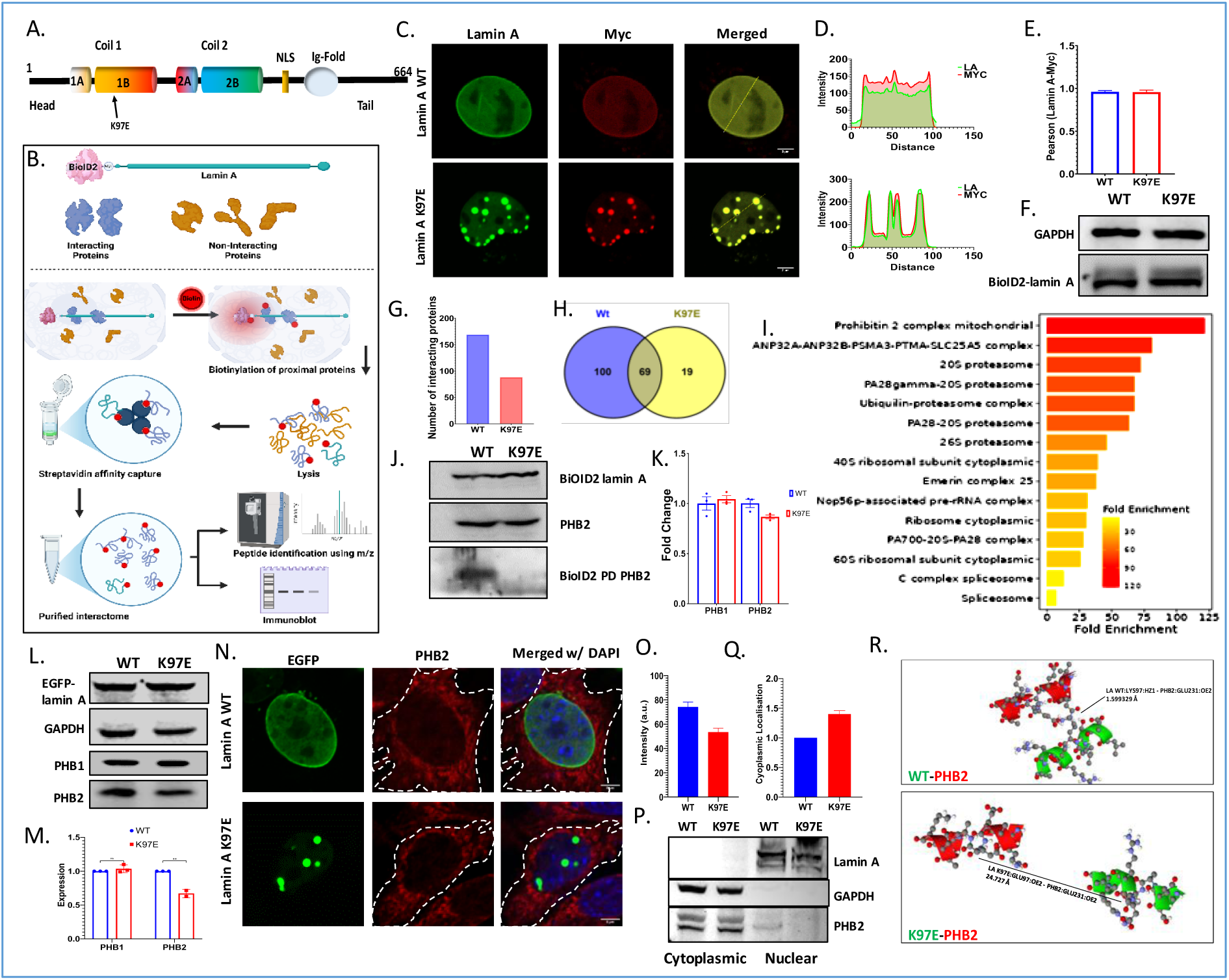
Loss of interaction with PHB2 complex in lamin A mutant K97E. A. Schematic representation of the K97E mutation in the 1B Domain of Lamin A (Not to scale). B. Schematic overview of proximity-dependent biotinylation and streptavidin pulldown (BioID2) strategy. C. Z-projected immunofluorescence images showing localisation of myc-BioID2-lamin A variants (scale: 2 μm). D. Intensity profiles across the nuclear axis depicting myc-BioID2-lamin A and overall lamin A distribution. E. Pearson correlation coefficient analysis showing colocalization between myc and lamin A signals. F. Immunoblot of lamin A/C and GAPDH showing equal BioID2 input loading and comparable expression of BioID2–lamin A variants in transfected HEK293FT cells. G. Number of interacting partners of WT and K97E lamin A as observed from protein identification through BioID2-pulldown followed by MS. H. Venn-diagram indicating independent and overlapping interactome of WT and K97E lamin A. I. Enrichment analysis of protein–protein interactions lost in the K97E mutant compared to WT lamin A. J. BioID2-based immunoblot analysis showing reduced interaction between lamin A and PHB2 in the K97E mutant compared to WT. K. Quantitative PCR analysis of PHB1 and PHB2 mRNA levels, revealing differential expression in response to the K97E mutation in transfected C2C12 cells. L. Immunoblot analysis of PHB1, PHB2, GAPDH, and lamin A in C2C12 cells. M. Densitometric quantification of PHB1 and PHB2 immunoblots, normalised to GAPDH. N. Z-projected immunofluorescence images of PHB2 showing altered subcellular distribution upon K97E lamin A expression (scale: 4 μm). O. Quantitative analysis of PHB2 fluorescence intensity from immunofluorescence imaging. P. Subcellular fractionation of cells transfected with - WT and K97E lamin A variants, followed by immunoblot against GAPDH (Cytosolic marker), lamin A (Nuclear marker) and PHB2. Q. Densitometric quantification of PHB2 levels in the cytoplasmic fraction, normalised to GAPDH. R. In-silico molecular docking of PHB2 with lamin A variants shows reduced interaction affinity with the K97E mutant, characterised by increased spatial separation of PHB2 from the mutation site.

### K97E lamin A alters mitochondrial dynamics via PHB2 and actin cytoskeleton remodelling

It is also established that PHB2 – the inner mitochondrial membrane protein interacts with the autophagosomal membrane-associated LC3 protein through an LC3-interacting region (LIR) upon mitochondrial depolarisation. So we focused, at this point, on mitochondrial homeostasis and how it might be adversely affected by PHB2. Abnormalities in mitochondrial dynamics are major hallmarks of DCM, leading to long-term ailment of tissues/organs having large energy demands, such as muscles and specifically the heart (*43*). The mitochondrial fission-fusion dynamics are tightly regulated by several pathways involving prohibitins, which act as central proteins to regulate this balance (Fig. 2 A). Taking cues from the previous result, we followed the trail to investigate whether the alteration of the PHB2 axis would affect the morphology and dynamics of the mitochondria adversely. Using confocal and Structured Illumination Microscopy (SIM) on transfected C2C12 myoblasts, we observed reduced mitochondrial length in the case of K97E compared to the WT (Fig. 2 B, C). We performed experiments to precisely calculate the mitochondrial fission-fusion rate to account for the mechanism of length shortening through time-lapse imaging of mitochondria tagged with CMXRos, followed by analysis using Mitometer (*44*). It is worthwhile to mention that mitochondrial fragmentation is one of the key signs of cellular stress (*45*–*47*). We observed an increased rate of fission with a concomitant decrease in the fusion rate in the case of the mutant protein (Fig. 2 D, E), indicating that the fission-fusion balance was perturbed. Due to this reduced size of the mitochondria, we also observed a sharp increase in mitochondrial dynamics through an elevated mitochondrial velocity (Fig. 2 F). To validate the claim that the fission-fusion balance was indeed perturbed, we checked the expression profile of two major fusion and fission proteins, Opa1 and Drp1, respectively. Quantitative PCR analysis revealed a reduced Opa1 expression but no significant increase of Drp1 (Fig. 2 G). Subsequently, we quantified the corresponding protein levels through immunoblot, which also elucidated a significant reduction in Opa1 levels while Drp1 remains much the same (Fig. 2 H, I). Furthermore, it is interesting to note that mitochondrial fusion protein Opa1, through a concerted post-translational processing and functionalization mediated by PHB complex, helps in mitochondrial fusion through inner membrane fusion and is closely associated with the pathogenesis of DCM (*48*). Hence, its downregulation and corresponding reduction in mitochondria fusion are justified by PHB2 depletion as reported earlier. Although it was previously reported that irrespective of Drp1 levels in cells, downregulation of Opa1 is enough to reduce mitochondrial length (*49, 50*), that does not completely explain the increased mitochondrial fusion rate with sufficient clarity in this scenario. Plagued by this apparent anomaly at this point, we turned to another important player in fission - mitochondrial associated actins (MAA), which not only regulate mitochondrial transportation for proper mitochondrial distribution and aids in fusion, but also play a vital role in mitochondrial fission through the formation of an actin ‘ring’ in ER-associated mitochondrial fission sites (*51*). Upon checking the distribution of MAA, we found out that indeed, the cells harbouring the laminopathic mutant showed an increased association of mitochondria and actin inside cells (Fig. 2 J, K), thus explaining the conundrum. Interestingly, Inf2, one of the actin regulators which, in tandem with the endoplasmic reticulum, mediate actin ring formation during mitochondrial fission (*51*), was also overexpressed in K97E (Fig. 2 L), thereby strengthening the probability of elevated fission events.

**Figure 2.**
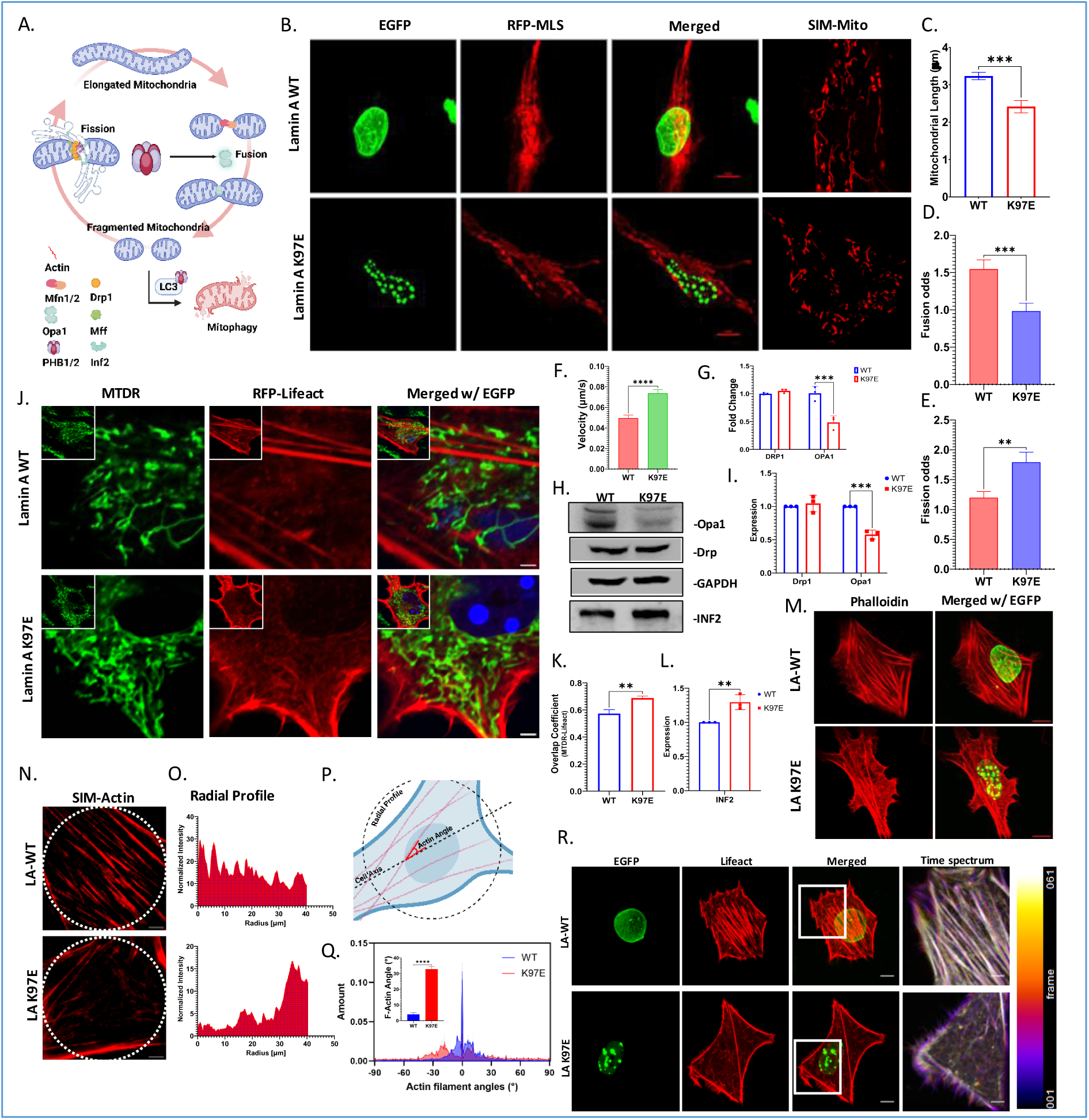
PHB2 mediated mitochondrial anomalies and role of actin destabilisation. A. Schematic representation of mitochondrial fission-fusion cycle highlighting the role of PHB2 (Not to scale). B. Z-projected confocal images of mitochondrial morphology visualised with RFP-MLS; super-resolution (SIM) images in the fourth panel reveal distinct mitochondrial morphology (scale: 5 μm). C. Quantified mitochondrial length from SIM images. D. Quantified mitochondrial fusion odds via time-lapse imaging of CMXRos-tagged mitochondria in lamin A-transfected cells. E. Quantified mitochondrial fission odds. F. Quantified mitochondrial velocity. G. qPCR analysis of Drp1 and Opa1 mRNA levels, revealing differential expression in response to the K97E mutation. H. Immunoblot analysis of Opa1, Drp1, GAPDH, and INF2. I. Densitometric quantification of Drp1 and Opa1 immunoblots, normalised to GAPDH. J. Live-cell confocal fluorescence imaging showing colocalization of RFP-lifeact–tagged actin filaments with Mitotracker deep red (MTDR)– labelled mitochondria in transfected cells; arrows indicate regions of colocalization (scale: 1 μm). K. Pearson correlation coefficient analysis showing colocalization between MTDR and RFP-lifeact signals in co-transfected cells. L. Densitometric quantification of INF2 immunoblots, normalised to GAPDH. M. Z-projected confocal images showing morphology and architecture of phalloidin-tagged actin (scale: 10 μm). N. Z-projected SIM images of phalloidin-labelled actin filaments; dotted circles indicate regions of interest (ROI) used for radial profile quantification (scale: 5 μm). O. Radial distribution profile of actin filaments quantified from a defined ROI. P. Representation of the quantification scheme for radial profile and actin filament angles. Q. Quantified distribution of actin filament angles; inset graph depicts mean actin angles. R. Time-lapse imaging of cells co-transfected with lamin A variants and RFP-lifeact; the first three panels represent Z-projected images for overall actin distribution (scale: 10 μm), while the fourth panel depicts the confocal plane time spectrum of actin dynamics within the marked ROI (scale: 5 μm).

This led us to investigate the actin and mitochondrial dynamics. We surmised that actin dynamics and/or polymerisation were modulating the mitochondrial fission-fusion equilibrium as a sequel to the mutation K97E. We observed through phalloidin staining that cells harbouring the lamin A K97E displayed disrupted filamentous-actin (F-actin) architecture as seen (Fig. 2 M). To obtain a clear perspective of the events, we employed SIM to scrutinise the F-actin filaments (Fig. 2 N), and observed that the actin filaments bundled more prominently in the cortex of the cell while displaying discontinuous/broken filaments throughout the cytoplasm as inferred from the radial profile (Fig. 2 O). The angle that the axis of the actin filaments subtends to the major axis of the cell (Fig. 2 P) exhibited an anisotropic distribution with higher values compared to the wild-type scenario (Fig. 2 Q), thereby implying a possible perturbation in cellular mechanical tension. Time-lapse imaging using RFP-lifeact showed that the actin filaments were highly dynamic or less stable in the mutant transfected cells (Fig. 2 R). In summary, the destabilised actin filaments could be a potential modulator of mitochondrial homeostasis. A lowered actin stress filament with increased dynamic of F-actin arises from intense cellular stress and is an indication of destabilised actins, which have been reported to directly affect mitochondrial morphology and functioning (*52*–*54*). Although it is now clear that the destabilised actins were indeed affecting the mitochondrial dynamics in K97E-transfected cells, a clear link between the lamin A mutation and actin destabilisation as a sequel to that remained largely elusive.

### Mechanistic insights into actin cytoskeletal dysregulation induced by K97E mutation

To assess the origin of severed actin filaments in the mutant transfected cells, we conducted F/G actin fractionation (Fig. 3 A) and found that the F/G actin ratio or the polymerised actin level indeed decreased in the mutant scenario (Fig. 3 B). To further elucidate the mechanism behind the decreased f-actins, we examined the proteins responsible for depolymerising actin filaments, notably cofilin. At first, we observed a straight downregulation of cofilin through checking its gene expression levels (Fig. 3 C), which was counterintuitive. Thus, we took a step back in its respective pathway to check the levels of ERK1/2, a previously known lamin A-regulated actin destabiliser which activates both cofilin and Arp2/3 complex (*55, 56*), and mediates actin depolymerisation through cofilin activation. Upon checking gene expression of ERK1, we observed a lowered mRNA level in K97E (Fig. 3 D). Further, we checked the expression of activated or pERK1/2 and found that the amount of pERK1/2 is also lowered in K97E-containing cells (Fig. 3 E, F). Upon confocal imaging of pERK1/2 (Fig. 3 G), we observed an expected lowered overall intensity with the majority of nuclear pERK sequestered in K97E aggregates (Fig. 3 H, I). This phenomenon might preclude ERK1/2 from activating cofilin and Arp2/3. Thus, combining reduced activated ERK1/2 levels in cells with cofilin downregulation indicates that depolymerisation of actins might be lowered in K97E and might not be the cause of reduced f-actins. But then, why is there more G-actin present in the mutant transfected cells?

**Figure 3.**
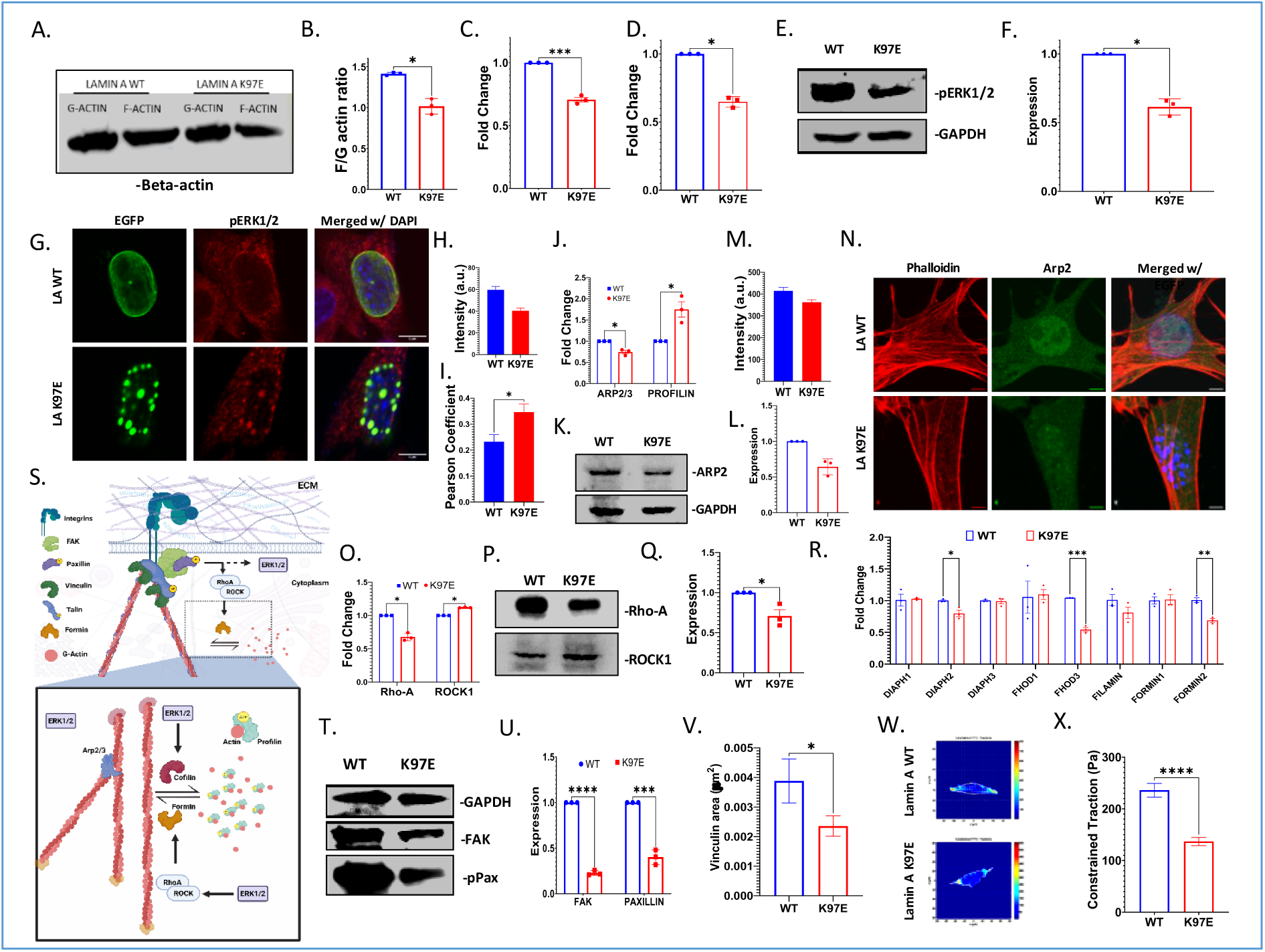
Actin destabilisation through the RHO-ERK-FAK axis also leads to defects in cellular mechanotransduction. A. Actin fractionation followed by immunoblot to quantify globular (g) and filamentous (f) actins. B. Quantification of F/G actin ratio from actin fractionation. C. qPCR analysis to check the mRNA levels of cofilin1. D. qPCR analysis to check the mRNA levels of ERK1. E. Immunoblot against phospho-ERK1/2 with respect to GAPDH. F. Densitometric quantification of pERK1/2 from immunoblots, normalised to GAPDH. G. Z-projected immunofluorescence images showing localisation of pERK1/2 in transfected C2C12 cells (scale: 5 μm). H. Quantitative analysis of pERK1/2 fluorescence intensity from immunofluorescence imaging. I. Pearson correlation coefficient analysis showing colocalization between pERK1/2 and lamin A variant signals. J. qPCR analysis of Arp2/3 and Profilin mRNA levels, revealing differential expression in response to the K97E mutation. K. Immunoblot against Arp2 with respect to GAPDH. L. Densitometric quantification of Arp2 from immunoblots, normalised to GAPDH. M. Z-projected immunofluorescence images of Arp2 showing subcellular distribution upon K97E lamin A expression (scale: 5 μm). O. qPCR analysis of RhoA and ROCK1 mRNA levels, revealing differential expressions in response to the K97E mutation. P. Immunoblot against RhoA and ROCK1. Q. Densitometric quantification of RhoA from immunoblots, normalised to GAPDH. R. qPCR analysis of mRNA levels of various actin polymerisation-promoting proteins, including formins, revealing differential expression in response to the K97E mutation. S. Schematic representation of focal adhesion, highlighting its role in actin polymerisation cycle (Not to scale). T. Immunoblot against Fak and phospho-paxillin with respect to GAPDH. Q. Densitometric quantification of FAK and pPaxillin from immunoblots, normalised to GAPDH. V. Quantification of vinculin puncta area marking focal adhesion complex. W. Representative images from traction force microscopy showing force distribution in cells. X. Quantified constrained traction forces exerted by cells.

Investigating further, we assessed the expression of Arp2 and profilin. Arp2 is an actin nucleator, which localises throughout the cell at actin branching sites, whereas profilin is a G-actin binding protein, which helps in initiating actin polymerisation through ATP-hydrolysis (*57*). Upon examining the gene expression of Arp2, we noted a slight downregulation, which was also reflected in protein expression levels in the case of K97E (Fig. 3 J-L), which, along with downregulated pERK1/2, suggests that the branching of actins may not be influencing actin dynamics significantly. This was validated by confocal images showing similar downregulation, particularly inside the nucleus (Fig. 3 M, N). Conversely, profilin exhibited a significant upregulation of gene expression (Fig. 3 J), which, in tandem with the previous observations, accounted for a larger pool of G-actin present in K97E-transfected cells compared to wild types. Subsequently, we checked the expression of RhoA and ROCK1, two major regulatory entities of actin polymerisation as well as a mitochondrial health determinant (*58, 59*). Gene expression supplemented by protein expression profile revealed a downregulation of RhoA (Fig. 3 O-Q), which is crucial for the activation of ROCK1 and, consequently, actin polymerisation through formins. Following RhoA downregulation, we checked the expression of several groups of formins and noticed that FHOD3, formin2 and DIAPH2 were downregulated in K97E (Fig. 3 R). Thus, we can summarise that it was due to lowered actin polymerisation and not increased which led to an increased G-actin pool in the cells.

Both RhoA and ERK pathways are interconnected with a third entity - FAK, which is also affected by lamins and formins like DIAPHs and FHODs (*60*). FAK is an integral part of the focal adhesion complex and interacts with both ERKs and RhoA in mechanotransduction (*61*–*63*), where it phosphorylates paxillin to initiate the signalling cascade (*64*–*66*). Along the same FAK also partakes in actin polymerisation through feedback from mechanical signals received from focal adhesion (*67*) (Fig. 3 S). While checking the expression of FAK and p-paxillin, we found out that both the proteins were severely depleted in K97E induced cells (Fig. 3 T, U), which was reported earlier to cause gross actin destabilisation and mechanotransduction defects (*68*). Depletion of FAK also indicates lowered focal adhesion stability, which, when checked through staining vinculin (Supplementary Fig. 3), elucidated a lowered vinculin area per focal adhesion (Fig. 3 V), amounting to weakened focal adhesions in K97E containing cells. Along with its role in mitochondrial homeostasis, actins play a primary role in cellular mechanotransduction. While it transmits the mechanical signals from focal adhesions to the nucleus via LINC complex (*60, 66*), it is also responsible for maintaining cellular stiffness and the generation of traction forces. Following observation on gross defects in actins and focal adhesion, we hypothesised mechanotransduction defects via alteration of traction forces exerted by the cell. Thus, to confirm that we employed traction force microscopy (TFM), and observed that the K97E cells indeed generated ∼50% lower traction forces compared to wild-type cells (Fig. 3 W, X).

In summary, we concluded that actin destabilisation is caused by a lack of proper actin polymerisation through downregulation RhoA-ERK-FAK axis in the cell, which not only contributes towards increased mitochondrial fusion but also a defective mechanotransduction pathway. Along the same lines, it must be mentioned that PHB2 is one of the proteins that also helps in the stabilisation and activation of FAK through the Akt pathway, along with its interaction with ERK and RhoA to maintain muscle homeostasis (*69*). Thus, through PHB2, we can correlate both actin and mitochondrial defects in cells having the K97E lamin A mutation.

### K97E mutation disrupts mitochondrial redox homeostasis and metabolic output

After concluding from our previous results that mitochondrial destabilisation is mediated largely by PHB2 expression and perturbations in actin polymerisation, we sought to investigate its functional implications. By and large, mitochondrial homeostasis correlates with its membrane potential (Ψ_m_), which is crucial for the functioning of mitochondrial enzymes and subsequent OxPhos pathways (*70*). The increased mitochondrial fragmentation often points towards unhealthy mitochondria, characterised by mitophagy and reduction of Ψ_m_ (*46*). To check the mitochondrial membrane potential, we used an indirect analysis by measuring CMXRos (*71*), which can only effectively stain the healthy mitochondria, having a differential membrane potential, leaving the damaged mitochondria unstained. On the other hand, MLS-RFP is a direct readout for both healthy and unhealthy mitochondria. Therefore, comparison of the mitochondrial footprints from MLS-RFP and those of CMXRos yielded an approximate measure of unhealthy mitochondria. We observed a significant reduction of Ψ_m_ of mitochondria in K97E-transfected cells (Fig. 4 A, B). Along the same line, we also noticed that the mitochondrial length was further shortened while stained with CMXRos (Fig. 4 C), as compared to MLS-RFP, in K97E. Thus, we can infer that both the mitochondrial length and Ψ_m_ decreased in the case of the mutant.

**Figure 4.**
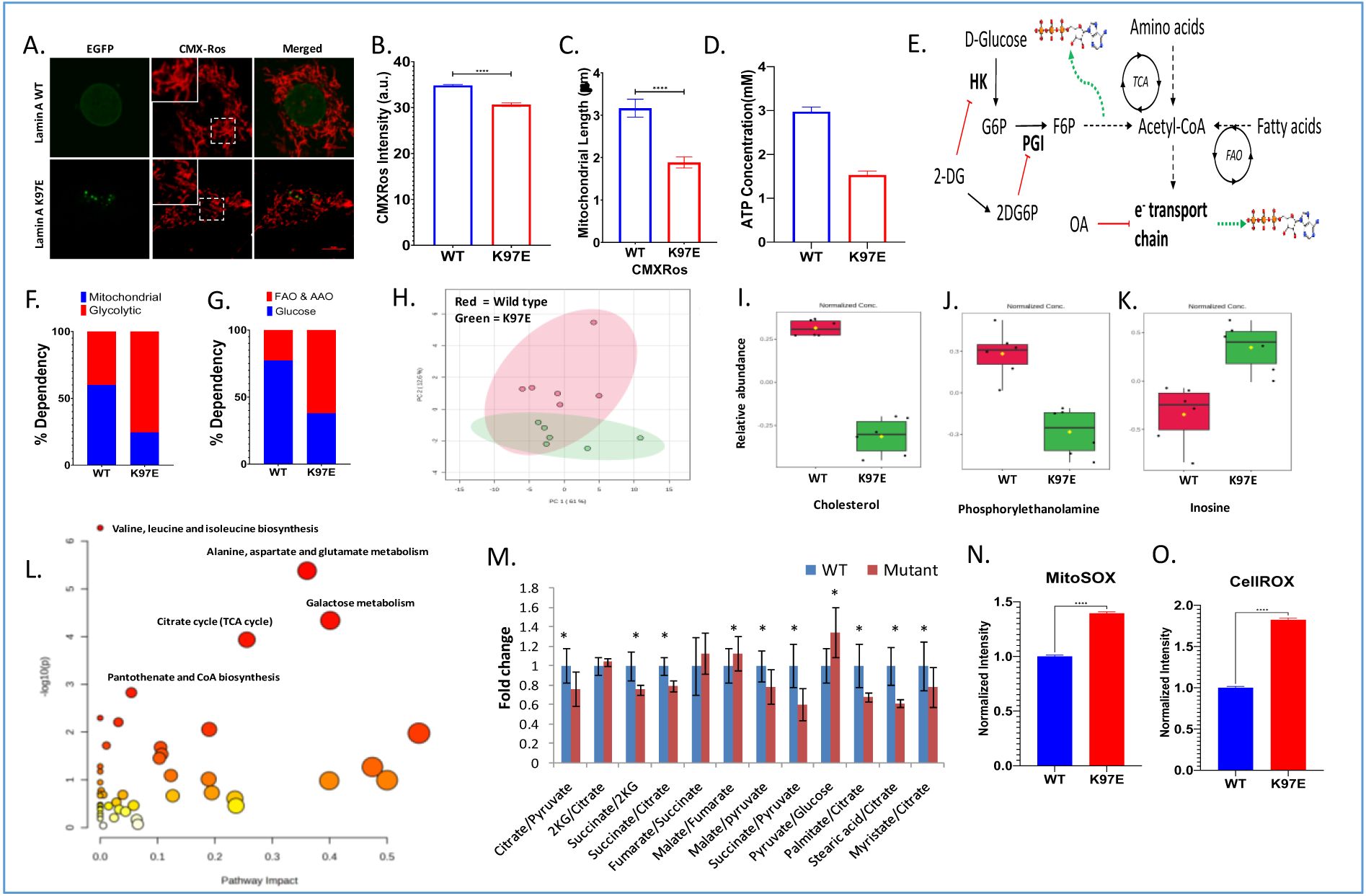
Altered metabolic profile leads to decreased ATP levels and elevated ROS. A. Confocal imaging of mitochondrial health visualised with CMXRos (scale: 10 μm). B. Quantitative analysis of CMXRos fluorescence intensity from confocal images. C. Quantified mitochondrial length from CMXRos labelled mitochondria. D. Quantified ATP concentration of cells transfected with lamin A variants. E. Schematic representation of the ATP dependency assay using 2-deoxyglucose (2-DG) and oligomycin (OA). F. Quantified mitochondrial vs glycolytic dependency of transfected cells. G. Quantified glucose vs beta oxidation, i.e. fatty acid oxidation (FAO) and amino acid oxidation (AAO) dependency of transfected cells. H. Unsupervised principal component analysis (PCA) of the K97E and WT metabolome. I-K. Relative abundance of cholesterol (I), Phosphorylethanolamine (J) and Inosine (K) obtained from metabolome analysis of transfected cells through GC-MS. L. Metabolic pathway enrichment conducted using significantly differentially abundant metabolites. M. Quantified ratio of selected metabolites from the bioenergetic pathway. N. Quantification of mitochondrial reactive oxygen species (ROS) using mitoSOX. O. Quantification of cellular ROS using cellROX.

To check the bioenergetic footprint of the K97E-containing cells, we measured the amount of ATP in the cells. A drastic reduction in ATP content was observed in cells expressing the lamin A K97E mutant (Fig. 4 D). At this point, it is important to delineate the contribution of mitochondrial and non-mitochondrial biochemical pathways leading to ATP depletion. To that end, we used selective pathway inhibitors like Oligomycin (OA) and 2-deoxyglucose (2-DG) as described protocol and depicted in Fig. 4 E and Supplementary Fig. 4 (*72*). Results showed that concomitant with an overall decrease in ATP, there was a significant (∼3-fold) decrease in mitochondrial contribution to ATP production upon K97E mutation vis-à-vis the WT (Fig. 4 F). Further, while dependency on glucose oxidation decreased, the contribution of fatty acid oxidation (FAO) and amino acid oxidation (AAO) increased significantly in the mutant (Fig. 4 G). In order to gain further insight into the metabolic alterations associated with expression of the K97E mutant, untargeted analysis of the cellular metabolome was performed. The unsupervised principal components analysis showed an inherent tendency of segregation of the K97E and WT metabolome (Fig. 4 H). The supervised orthogonal projection to latent structures-discriminant analysis (OPLS-DA) of the metabolomic signature showed clear separation of the WT and K97E metabolome (Supplementary Fig. 5 A). The permutation test (100 permutations) showed a p value < 0.05, indicating goodness of model fit and predictive accuracy (Supplementary Fig. 5 B). Among metabolites that contributed to the separation of metabolome, cholesterol (Fig. 4 I), phosphorylethanolamine (Fig. 4 J) were found to be notably depleted, whereas inosine (Fig. 4 K) was found to be elevated in the K97E mutant even after FRD correction. Metabolic pathway analysis was conducted using differentially abundant metabolites (fold change > 1.25, P< 0.05) identified by volcano plot analysis (Fig. 4 L). It indicated a significant alteration in multiple pathways as listed in Supplementary Table 2. Particularly, alterations in TCA cycle, valine, leucine and isoleucine metabolism, alanine, aspartate and glutamate metabolism were significant even after FDR correction. In addition, the pantothenate and CoA biosynthesis pathway, which is essential for entry of glucose-derived pyruvate into the TCA cycle as well as metabolism of fatty acids and branched-chain amino acids, was also altered. Mitochondria play an important role in the metabolism of these amino acids and fatty acids via the TCA cycle. Taken together, these points to a coordinate derangement of mitochondrial metabolism as was indicated by the energetics assay. In a biochemical pathway, the ratio of two interconnected metabolites has been shown to offer a surrogate signature for biochemical flux and the relative enzymatic activities through the steps in between (*73*–*77*). Hence, we explored the ratio of metabolites involved in energy-producing metabolic pathways to get a more detailed picture of the effect on the energy metabolism. As shown in Fig. 4 M, there was a decrease in citrate/pyruvate, succinate/pyruvate, malate/pyruvate, succinate/2-ketoglutarate and succinate/citrate ratios in the mutant, indicating a low TCA cycle activity. This was also associated with an increase in the pyruvate/glucose ratio in the mutant, plausibly due to a lack of glucose-derived pyruvate utilisation. Therefore, an impaired glucose metabolism towards ATP production was also indicated by a decrease in glucose dependency in the mutant in the ATP assay. On the other hand, there was an increase in the gluconic acid/glucose ratio (Supplementary Fig. 5 C) in the mutant, indicating oxidative stress. Interestingly, the ratio of palmitate to citrate, as well as myristate or stearate to citrate, was significantly lower in the mutant, which may be a result of elevated FAO, as was indicated by the ATP assay and/or a decrease in *de novo* fatty acid synthesis due to low ATP availability. This is also reflected in the lower AMP/adenine ratio along with a lower cholesterol to citrate ratio (Supplementary Fig. 5 C), further strengthening the FAO over *de novo* fatty acid synthesis. Subsequent analysis of the amount of superoxide indicated an elevated oxidative stress as a sequel to metabolic abnormalities. To validate this further, we measured cellular and mitochondrial superoxide levels and observed that both of these superoxide species in K97E-transfected cells were increased, indicating an increased superoxide generation (Fig. 4 N, O) due to the elevated FAO.

## Discussion

The pathogenesis of DCM in the context of lamin A mutations has traditionally been attributed to two primary mechanisms: mechanical disruption of the nuclear envelope and aberrant gene regulation through altered chromatin-lamina interactions. In this study, we propose an additional mechanistic axis initiated by the K97E mutation in lamin A that disrupts mitochondrial and cytoskeletal homeostasis through destabilisation of PHB2 function. Our integrated model highlights PHB2 as a nodal factor linking nuclear and mitochondrial homeostasis, with broad implications for understanding DCM pathogenesis and guiding therapeutic strategies.

Lamin A constitutes a smooth, peripheral lamina essential for nuclear integrity and chromatin organisation. Our qualitative BioID2 based interactomics study identified that lamin A interacts with major protein groups including *PKM, ENO1, PGK1, TPI1, ALDOA, TKT, TALDO1,* and others, enriching towards ATP Synthesis, Pentose phosphate pathway, Glycolysis, and other carbon metabolism pathways (Supplementary Figure 6 A, B), whose presence in nucleus is well reported and is essential for nuclear bioenergetics and epigenetic remodeling (*78*–*81*). These interactions suggest a previously underappreciated role of lamin A in nuclear metabolic regulation, beyond its established structural functions. Although the functional relevance of these interactions remains to be elucidated. The K97E mutation of lamin A disrupts its filament assembly, and by extension, lamina formation.

Our recent study demonstrates that this accompanies the phenomenon of liquid-liquid phase separation of K97E mutant proteins due to increased self-association and disrupts lamin-heterochromatin association, suggesting a novel phase-separation behaviour not previously described in intermediate filament proteins (*82*). This could also explain the reduced interactome of the K97E mutant, as observed in our BioID2-MS dataset, enriching towards the PHB2 complex. Along the same lines, reciprocal IP, and molecular docking converge on a model in which lamin A interaction with PHB2 is abolished, depleting the lamin A-PHB2 complex in the nucleus, and enriching PHB2 towards the cytoplasmic fraction of the cell along with its overall downregulation (Fig. 5). PHB2 is not only limited to its role in mitochondrial homeostasis but also plays key role in several cellular processes that are often overseen (*41, 69*). Previously, PHB2 has shown a strong correlation with the pathogenesis of DCM, as it was observed that its depletion can lead to a distinct shift in the OxPhos pathway and reduced ATP levels, leading to heart failure (*25*). This is in association with PHB2’s interaction with DNAJC19, a mitochondrial chaperone, whose alteration is directly correlated to the pathogenesis of DCM with ataxia (*83*). Concomitant with PHB2 depletion, mitochondria in K97E cells display significant fragmentation as seen through shortened branch lengths, and increased motility arising from an increase in fission events accompanied by a decrease in fusion frequency, in accordance with previous reports (*25, 84*). Further, we revealed that PHB2 mediated downregulation of Opa1 leads towards a decreased mitochondrial fusion. Similar to PHB2, Opa1 is another member of the mitochondrial proteins, which shows a strong correlation with the pathogenesis of lamin A-mediated DCM. It has been reported that mitochondrial fragmentation due to improper Opa1 processing, in which PHB2 also plays a role (*23*), might lead to heart failure in mice (*85*). Additionally, elevated E2F/TP53 DNA damage response activity during lamin A mutation has also been reported to destabilise Opa1, leading to the development of DCM (*48, 86*). Examination of γH2AX levels in K97E-expressing nuclei revealed a marked increase in nuclear pH2AX foci (Supplementary Fig. 7 A-C), indicative of elevated DNA damage. This genomic instability may stem from increased ROS levels or from impaired DNA repair capacity due to the loss of functional lamin A, which is known to play a critical role in the maintenance of genome integrity (*87*). Such DNA damage may further contribute to the destabilisation of Opa1, as previously implicated through E2F/TP53-mediated pathways. Further, we explained the increased mitochondrial fission axis through profound actin cytoskeletal disruption in K97E-transfected cells. We observed disorganised actin structures, consistent with impaired Rho-ERK-FAK signalling and reduced formin-mediated polymerisation (Fig. 5). The surplus G-actin fuels the mitochondrial “actin cloud”, enhances the ER-mitochondrial contacts and recruits DRP1 to promote mitochondrial fission (*88*). Here, we can also appreciate that the nuclear exclusion of PHB2 can also impair transcriptional programs vital for cardiomyocyte survival, exacerbating disease progression (*69*). As PHB2 is a transcription factor for both ERK1 and 2 (*89*), and its nuclear exclusion might trigger the downregulation of the MAPK pathway in K97E lamin A mutation, leading to a destabilised actin fraction. As mitochondrial health declines, two simultaneous, yet associative phenomena occur, which directly lead to cytoskeletal, to be precise, actin remodelling. First is ATP depletion, without which profilin-actin-ATP complex cannot be attained, and formin-mediated polymerisation could not take place (*90*). Secondly, the generation of cellular ROS is largely associated with cytoskeletal remodelling and cortical actin strengthening in cells, as observed in our system (*91, 92*). In either case, the associated actins lose their integrity and mediate abnormalities in mechanotransduction signals. In our case, we observed altered traction forces exerted by cells, which is a key inside-out mechanical signal that helps cells communicate and create an optimal niche for tissue growth. Without a proper mechanical crosstalk between cells, we can expect a dysregulation of several cellular processes, such as abnormal cell division or improper differentiation. As muscle myoblasts heavily rely on ATP and actin stress fibres for optimal cell differentiation (*93, 94*), one can expect an alteration in this vital process. Studies expanded over the past decade already showed that the cardiomyopathy pathogenesis is frequently linked to underlying metabolic disturbances (*95, 96*), and the pathogenesis of lamin A-mediated DCM can be further explored through the functional analysis of lamin’s role in metabolic regulation. Multiple interactions between the nuclear lamina and fatty acid metabolism have been reported earlier (*97*–*99*), along with the existence of lamin-mediated lipodystrophies, but their exact pathogenesis of DCM associated with A-type lamin defects has not been proven until very recently (*100*). Our untargeted metabolomics uncovered dysregulation in fatty acid beta-oxidation, amino acid biosynthesis, and several bioenergetic pathways in K97E-transfected cells, indicating a global collapse of metabolic homeostasis leading to an elevated ROS level. Although it has been observed on a common basis that optimised FAO is very crucial for cardiac maintenance and provides the majority of the energy sources required for the cardiac tissues to function, while maintaining ROS. But depleted PHB2 levels are highly associated with an improper and incomplete FAO in cardiac cells, leading to elevated ROS and depleted ATP levels in the cell, along with improper ROS management (*25, 101*). Along with elevated FAO, several other metabolites are also observed to change, among which Inosine is seen to get upregulated in K97E-transfected cells. Inosine was shown to increase mitochondrial biogenesis and mitophagy as well as mitochondrial respiration through OxPhos by eliciting TCA cycle enzymes, which could be an adaptive response for survival (*102, 103*).

Our investigation into lamin-associated cardiomyopathy, conducted using a transient cellular model in the absence of in vivo or patient-derived systems, naturally invites scrutiny regarding the translational validity of our findings. However, the extensive and well-established links between lamin A and metabolic regulation substantiate our conclusions. Lamin A-mediated FPLD, MAFLD, lipid metabolism abnormalities in MAD(*98*), regulation of mTOR signalling cascade, which is repeatedly seen to get disrupted during laminopathies (*104*) and interplay with SIRT1 (*33*) have repeatedly linked lamin A with metabolism. Along the same lines, the dysregulation of LADs during lamin A mutation is also seen to regulate the metabolic fate of the patient (*105*). Similarly, *LMNA* knockout mice mimicking DCM are also reported to show impaired complete β-oxidation through accumulation of medium and long chain acylcarnitines along with depleted high-energy nucleotides such as ATP, ADP and AMP (*106*). The most notable phenotypic parallel of our mechanistic framework came from a very recent patient study which showed lamin A variant G105L, located on the same domain as K97E and one of the founder lamin A mutation leading to DCM showed similar trends of reduced mitochondrial heath, biomass and oxygen consumption rate, reduced ATP production along with elevated ROS (*107*). Thus, in cardiac cells, where demand for ATP and mitochondrial health is paramount-PHB2 dysregulation and the resulting mitochondrial dysfunction looped with actin abnormalities likely drive contractile deficits and arrhythmogenesis. Our work uncovers a previously unrecognised pathway by which lamin-A mutations perturb PHB2 homeostasis, triggering cytoskeletal remodelling and mitochondrial dysfunction that converge on bioenergetic collapse and genomic stress. By integrating nuclear architecture, cytoskeletal dynamics, and mitochondrial physiology, our study provides a framework for targeted interventions in laminopathies.

## Materials & Methods

### Subcloning in BioID2 plasmid

Lamin A wild-type and lamin A K97E were cloned as reported earlier (Bhattacharjee et al. 2013) in the pEGFP vector (Clontech). EGFP-lamin A variants were used as templates for PCR. The purified PCR was digested with EcoRI & BamHI and sub-cloned into BioID2-myc plasmid, received as a gift from Dr. Kyle Roux, in which we sub-cloned lamin A WT and K97E using the cloning primers:

Sense: CCGGAATTCCGGATGGAGACCCCGTCCCAG

Antisense: CCGGGATCCCGGTTACATGATGCTGCAGTTCT

The sub-cloned construct was sequenced and validated.

### Cell Culture and Transfection

C2C12 cells (ATCC) were cultured in ATCC-formulated DMEM supplemented with Penicillin-streptomycin and Glutamax (Gibco) as previously described(*37, 108, 109*). HEK293T (ATCC) cells were maintained using the same parameters as C2C12. The cells were maintained according to standard protocols at 37℃ and 5% CO2. Cells at passages 2 to 3 were seeded to a confluency of 50-60% for transfection. For each transfection, 1:1.5 of DNA with Lipofectamine 3000 (Invitrogen, USA) was utilised, following the manufacturer’s instructions. Cells were processed for subsequent experiments 24-36 hours post-transfection.

### Staining for immunofluorescence

Cells were fixed with 4% paraformaldehyde for 20 minutes at room temperature. Subsequently, the fixed cells were permeabilised with 0.5% Triton X-100 for 6 minutes and then incubated with primary antibodies diluted in 5% NGS in PBS for 2 hours at room temperature or overnight at 4°C (according to the manufacturer’s protocol). Following which, cells were washed thrice with 0.5% Tween-20 in PBS and thrice with PBS before incubating with fluorochrome-conjugated secondary antibodies for 1hour at 37°C. It was followed by washing steps. Finally, the cells were mounted with Vectashield (Fisher Scientific, USA).

### Fluorescence Microscopy and Image Analysis

For confocal imaging, the slides were visualised using a 60X water or 100X oil immersion objective with a numerical aperture of 1.4 and a refractive index of 1.515 on a NIKON TiE inverted research microscope (Nikon, Japan) equipped with a digital zoom, as described in previous literatures (*1*). For XY and Z-stack imaging, a resonant scanner with a line averaging of 4.0 was utilised. Z-stack images were acquired with a step size of 0.15 µm. Live cell imaging was performed with a culture dish under a thermal incubator (Tokai Hit) maintained at 37°C with cells in CO2-independent media.

For super-resolution structured illumination microscopy, a super-resolution Plan Apochromat TIRF 100x/1.5 NA/WD 0.13 mm objective (Nikon, Japan) was employed to capture the morphology and distribution of RFP-lifeact-decorated actins and MLS-RFP marked mitochondria within cells (*108, 110*).

Images were exported using NIS-Elements Analysis AR (Ver. 4.13), and ImageJ software (Ver. 1.8.0_112) was utilised for all confocal, super resolution and immunoblot image analysis. Mitochondrial length, fission-fusion rate and velocity were calculated using Mitometer using default parameters (*44*).

### Proximity Biotin labelling and mass spectrometric analysis

#### Sample Preparation

HEK293T cells were initially cultured up to the 3rd passage, when transfection was carried out with the plasmids in four 100 mm dishes using calcium (*111*). Following 24 hours of transfection, the media were replaced with complete media supplemented with 50μM of Biotin. The cells were then cultured for an additional 18 hours before harvesting. Before harvesting, cells were washed thrice with ice-cold PBS and lysed according to the protocol outlined by Kim et al(*112, 113*). Specifically, 2 ml of BioID lysis buffer containing 50 mM Tris (pH 7.4), 500 mM NaCl, 0.4% SDS, 1 mM dithiothreitol, and protease inhibitor cocktail (PIC) was added to the scraped cell pellet, homogenised, and sonicated for 3 minutes at 50% amplitude. Subsequently, 2% Triton X-100 was added to the lysate, followed by another round of sonication for 5 minutes. Finally, 2 ml of 50 mM Tris was added to the lysate, and a third round of sonication was performed for 5 minutes under ice-cold conditions. The lysate was then clarified by centrifugation at 13,000g for 10 minutes. The cleared lysate was then added to 30μl of Streptavidin agarose beads (Pierce) and incubated overnight at 4°C with shaking. The following day, the beads were washed sequentially with wash buffer 1 containing 2% SDS in 50 mMTris, wash buffer 2 containing 0.1% (w/v) deoxycholic acid, 1% (w/v) Triton X-100, 1 mM EDTA, 500 mM NaCl, and 50 mM HEPES (pH 7.5), wash buffer 3 containing 0.5% (w/v) deoxycholic acid, 0.5% (w/v) NP-40, 1 mM EDTA, 250 mM LiCl, and 10 mM Tris·Cl (pH 7.4), and finally with wash buffer containing only 50 mM Tris. The beads were then collected by centrifugation, and 100μl of protein loading buffer was added. The mixture was heated at 95°C for 15 minutes before analysis.

#### Mass spectrometric sample preparation and analysis

Protein samples were in-gel digested as described by Rawat et. al(*114*). Digested peptides were analysed on a Q-Exactive mass spectrometer coupled with a Nano flow HPLC system (Easy-nLC 1200 Thermo Scientific). The sample was loaded onto an Easy Spray Column PepMapTM RSLC C18 (3 µm, 100 A0, 75 µm x 15 cm). Samples were eluted with a 60 min gradient between solution A (5% acetonitrile, water containing 0.2% formic acid) and solution B (95% acetonitrile in water containing 0.2% formic acid) at a flow rate of 300 nl/min. Mass spectra were acquired with an Orbitrap analyser at the resolving power of 70,000 at m/z 200. The scan range selected was 400-1650 m/z. MS/MS was carried out in HCD (normalised collision energy - 30%) mode with a resolving power of 17,500 at m/z 200. TopN 10 ions were taken for MS/MS. The lock mass option was enabled for accurate mass measurements.

The raw data files were searched against the Protein Database: Human fasta database-5JAN-2023-HUMAN-uniprot-download_true_format_fasta_query 28homo_20sapiens_29_20AND_-023.01.05-06.28.17.74.fasta (20330 Sequences) & Contaminant FASTA database containing (298 Sequences) using Sequest HT search engine on Thermo Proteome Discoverer (Version 2.2.0388). N-terminal acetylation and methionine oxidation were included as dynamic modifications; carbamidomethyl of cysteine as a static modification. A Precursor Mass Tolerance of 10 ppm and a fragment Mass Tolerance of 0.05 Da were used. FDR setting was 0.01 & 0.05 for both peptide and protein levels; Percolator q-value validation; Delta Cn was 0.05. The following filters were applied to the PD analysed data: i. master is equal to master, ii. minimum 1 unique peptide & 1 peptide/protein, iii. high Peptide confidence.

#### Analysis of interactome

For differential analysis of interactome, Venny 2.0 was utilised after filtering the raw data for the proteins which are homogeneously highly abundant in the sample replicates. The loss of interaction (WT ∩ K97E’) was enriched using ShinyGO (ver 0.77) using the CORUM PPI database. Further, the interaction with PHB2 was confirmed using IP/IB.

### Immunoprecipitation

Transfected cells were washed with ice-cold PBS, scraped, and pelleted in 1.5 mL microfuge tubes. The pellets were stored at −80°C until further processing. For lysis, 1 mL of RIPA buffer (50 mM Tris-Cl pH 7.5, 150 mM NaCl, 1% NP-40, 0.1% SDS, 0.5% sodium deoxycholate, 1 mM EDTA, and 1× protease inhibitor cocktail) was added per 10⁷ cells (∼750 µL for a 90 mm dish), followed by sonication. The lysate was pre-cleared by incubating with 25 µL of equilibrated Dynabead Protein-G per mL of lysate at 4°C for 1 hour with rotation. Beads were then pelleted using DynaMag. The cleared lysate was divided into two equal portions: one incubated with the anti-myc antibody and the other with normal rabbit IgG (negative control). Both were incubated overnight at 4°C with 50 µL of Dynabead Protein-G (Invitrogen) per mL of lysate. Beads were washed thrice with lysis buffer. Bound proteins were eluted with 30 µL Laemmli sample buffer (5% β-mercaptoethanol) and heated at 95°C for 15 min. Eluates were analysed by immunoblotting.

### Immunoblot

Post-lysis protein concentration was assessed using the Bradford assay, following previously established protocols(*1, 37, 108*). Equal volumes of protein samples and Laemmli buffer were combined and boiled at 95°C for 10 minutes and cooled before gel loading. Subsequent to gel loading, proteins were separated via 10% SDS-PAGE and transferred onto a nitrocellulose membrane (Merck Millipore, USA). The Immunoblot procedures were conducted as described in prior literature(*87, 115*) with antibodies mentioned in Supplementary Methods Table 1. Band intensities were quantified using ImageJ.

### Quantitative PCR

Quantitative PCR was performed as per the previous standardised protocol by Sengupta et al.(*1*) with primers mentioned in Supplementary Methods Table 2.

### Nuclear Fractionation

A subcellular fractionation (SF) buffer was produced by dissolving 20 mM HEPES (pH 7.4), 10 mM KCl, 2 mM MgCl_2_, 1 mM EDTA, 1 mM dithiothreitol (DTT), 1 mM EGTA, and PIC. Cells grown on two 60mm culture dishes were washed thrice with ice-cold PBS and scraped into 400 µL of the cold fractionation buffer, followed by incubation on ice for 15 minutes. The cell suspension was then mechanically ruptured using a 27-gauge syringe, followed by another 20-minute incubation on ice. The lysate was then centrifuged at 720 ×*g* for 5 min to separate the nuclear pellet from the cytoplasmic supernatant, including mitochondria and membranes. The nuclear pellet was washed with 500 µL SF buffer, gently resuspended by pipetting, and homogenised further by passing through a 25-gauge needle to shave off impurities. The sample was centrifuged again at 700 ×*g* for 10 minutes, after which the supernatant was discarded, and the nuclear pellet was then lysed using RIPA and processed for IB.

### Homology Modelling and Molecular Docking

The 3D structure of the target protein was generated using homology modelling via SWISS-MODEL (https://swissmodel.expasy.org/), using 8J4I and 6JLB as templates and validated. For molecular docking, the modelled proteins were submitted to the HADDOCK web server (https://wenmr.science.uu.nl/haddock2.4) and rigid-body docking, followed by energy minimisation and water refinement, were performed to generate multiple complex conformations (https://doi.org/10.1038/s41596-024-01011-0). The top-ranked cluster, based on HADDOCK score and binding energies, was selected and analysed for binding efficiencies.

### Actin Fractionation

To separate G-Actin (globular actin) and F-Actin (filamentous actin), cells are first grown in a 60mm dish until they reach approximately 60% confluency. Following this, the cells are washed twice with ice-cold PBS. Subsequently, Actin Stabilising Lysis Buffer, composed of 0.1M PIPES (pH 6.9), 30% Glycerol, 5% DMSO, 1mM MgSO4, 1mM EGTA, 1% Triton X-100, 1mM ATP, and 1X PIC, is added, and the cells are incubated for 10 minutes. After scraping the cells and transferring them to a 1.5ml Eppendorf tube, centrifugation is performed at 10,000g for 75 minutes at 4°C. The resulting supernatant, containing G-Actin, is collected in a separate Eppendorf tube and mixed with an equal volume of 2X loading buffer. Meanwhile, the pellet, enriched with F-Actin, is dissolved in 400μl of 1X loading buffer. Following this, both fractions are heated for 10 minutes at 100°C to denature proteins. Finally, equal amounts of each fraction are loaded onto a gel, and separation is confirmed using immunoblotting.

### Traction Force Microscopy

#### Preparation of TFM Substrate

Traction force microscopy (TFM) utilising the bead tracking method is conducted on polyacrylamide substrates with a stiffness of 10 KPa. The preparation of these substrates follows the procedures outlined by Kulkarni et al.(*116*), with minor modifications. A 27mm glass bottom dish is initially treated with 10% methanolic APTES (Sigma, USA) for 15 minutes, followed by washing and subsequent treatment with a 0.5% glutaraldehyde solution (Sigma, USA) for 45 minutes. The dish is thoroughly washed, air dried, and immediately utilised. Concurrently, a polyacrylamide solution with a stiffness of ∼10 KPa is prepared according to the methodology described by Tse and Engler(*117*). This involves the addition of 10% acrylamide, 0.1% bis-acrylamide in water, along with 0.01% 0.2μm FluoSpheres (Invitrogen, USA), followed by thorough homogenization. Subsequently, the solution is polymerised using 10% APS and TEMED. Upon addition of the crosslinker and catalyst, 35μl of the mixture is poured into the silanated glass bottom dish. A poly-L-lysine-coated 22mm coverslip is then inverted onto the mixture to create a thin layer of polyacrylamide substrate. After 30 minutes, the coverslip is carefully removed, and the gel is washed to remove any excess acrylamide. Subsequently, 50mg/ml Sulpho-SANPAH is applied to cover the polymerised gel, followed by exposure to UV light for 30 minutes. After aspiration of the solution, 1ml of 100μg/ml fibronectin is pipetted onto the activated gel and incubated overnight at 4°C. Following the incubation period, the gel is washed thrice with PBS and sterilised using UV sterilisation before cell seeding.

#### Image Acquisition for TFM

Transfected cells are trypsinised after 24 hours of transfection and seeded into the prepared gels at a very low confluency (1 cell/100 μm) to ensure visibility of a single cell per frame. The cells are allowed to grow for at least 18 hours before image acquisition. 1ml of CO2-independent media is added to the dish to maintain uniform conditions throughout the experiment. Imaging is performed using a 40X objective on the Nikon inverted fluorescent microscope. Initially, a DIC image is captured along with the initial bead positions using a 561nm laser. Subsequently, 1ml of 10X (2.5%) trypsin is used to detach the cells for 30 minutes without disturbing the dish. After 30 minutes, another image is taken to capture localised bead movement. The images are then converted to TIFF format for subsequent traction force calculation.

#### Traction force measurement

The MATLAB algorithm developed by Kulkarni et al. is utilised for traction force measurement. The exact protocol outlined in their work is followed, employing the acquired images to obtain the constrained traction forces exerted by the cells.

### ATP dependency determination

ATP dependency was carried out by protocols followed by Islam et al.(*72*) with slight modifications. Cells were seeded in replicates in a white walled and clear 96-well plate with a seeding density of 10^4^ cells per well, followed by transfection. 24 hours post-transfection, the cells were supplemented with growth media containing 10 0mM 2-deoxyglucose (2-DG), 1μM oligomycin (OA) and a third set with a combination of both. 0.01% DMSO-treated cells were taken as a control. After an hour of incubation with the drugs, the cells from the white-walled plate were lysed for 15 minutes with constant shaking, followed by ATP determination using the ATP determination kit (Abcam) by quantifying the luminescence emitted. The data was normalised with the viability assay performed in the replicate clear plate using the MTT assay.

After which, the dependencies were calculated using the formulas:

- Glucose dependence (%) = [(DMSO control – 2-DG)/ (DMSO control – DGOA)] X 100.
- FAO & AAO capacity (%) = 100 – Glucose dependence.
- Mitochondrial dependence (%) = [(DMSO control – OA)/ (DMSO control – DGOA)] X 100.
- Glycolytic capacity (%) = 100 – Mitochondrial dependence.

### Cellular and mitochondrial superoxide determination

Cells were seeded in replicates in black walled and clear 96-well plates with a seeding density of 10^4^ cells per well, followed by transfection. 24 hours post-transfection, the cells were supplemented with 500 nM MitoSOX or 5 μM CellROX Deep Red (Invitrogen) and incubated at 37°C for 30 min. After which the cells were washed with PBS and the fluorescent emission was measured to obtain the relative superoxide levels.

### Metabolome analysis using MS

#### Sample Preparation

C2C12 cells were initially cultured until reaching the third passage, after which they were transfected with the plasmids in 6-well dishes using Lipofectamine 3000. Following 24 hours of transfection, cells were washed twice with 150 mM NaCl at the indicated time points before metabolite extraction. Metabolite extraction and analysis were performed following a modified version of the protocol described earlier (*118*). Briefly, 500 μl of extraction solvent (Water: Acetonitrile: Isopropanol = 2: 3: 3) containing 10 μM of 3-amino-4-methoxybenzoic acid as internal standard was added to the cell culture plate. Then, cells were scraped with cell scrapper and collected in 1.5 ml Eppendorf tubes and stored in −80 °C freezer until further processing. Cell lysis was achieved through three cycles of rapid freeze-thawing using liquid nitrogen and a 37 °C incubator, followed by centrifugation at 13,000 ×g for 30 minutes at 4 °C. A 100 μl aliquot of the supernatant was transferred into a 0.6 ml glass crimp-top microvial (27312, Supelco) and dried in a vacuum concentrator for approximately one and a half hours. Each sample was then treated with 30 μl of 2% methoxamine (MOX) reagent (TS-45950, ThermoScientific™, USA) and incubated at 50 °C for one hour with the lid closed to enable methyloxime derivatisation. After cooling to room temperature, the samples were silylated by adding 50 μl of MSTFA reagent (M-132, Supelco, USA) and heating at 65 °C for one hour in a sealed heat block. Blank samples were prepared using only the extraction solvent and following the same procedure. For quality control (QC), pooled samples were created by combining equal volumes of metabolite extract from all samples.

#### GC-MS analysis: Mass spec protocol

Samples were analysed with a 7890B GC coupled to a 5977B single-quadruple mass spectrometer (Agilent, USA). The chromatographic separation of analytes was achieved on an HP-5MS column (30 m × 0.25 mm × 0.25μm) with helium as the carrier gas. The front inlet was used in splitless mode at a temperature of 300°C. The oven temperature was maintained at 70°C for 5 minutes, followed by a ramp to 280°C at 5°C /minute. The temperature was then increased to 295°C at 10°C/minute and held at 295°C for 4 minutes. MS source and MS quad temperatures were set to 230°C and 150°C, respectively. The EI-MS spectra were acquired in full scan mode in the m/z range of 45-500. Samples were run in randomised order with intermittent injections of blank a nd pooled QC samples.

#### Data analysis

Column performance and consistency of instrument response were checked by manual inspection of chromatograms using MassHunter qualitative analysis software (Agilent, USA). Data was deconvoluted to extract features using MassHunter quantitative analysis software. Compound identity was based on the NIST library score as well as matching with authentic standards wherever available. Peak integration was performed with at least 2 unique ions for that feature. Features showing significant peaks in blank samples were removed. The blank areas were subtracted from the rest of the features. All features were normalised with respect to the internal standard as well as the protein concentration of each sample measured post-centrifugation during sample preparation. Features showing CV >30% in the pooled QC samples were removed from further analysis. The data were then sum-normalised, log-transformed and Pareto-scaled for multivariate analysis using MetaboAnalyst 5.0 (*119*) (https://www.metaboanalyst.ca/). Unsupervised PCA and supervised OPLS-DA analysis were employed to identify group segregation patterns as well as to find features contributing to the segregation. Volcano plot analysis was used to screen for features showing fold change in relative abundance > 1.5 (with raw p value <0.05) between the wild type and the mutant, which were used for analysis of metabolic pathways using MetaboAnalyst 5.0.

## Statistical analysis

All the statistical analysis was done using GraphPad Prism (Version 10). Unpaired Student T-test was used to determine the significance of the results with a 5% tolerance, unless mentioned. Immunoblotting and qPCR experiments were each performed in three biological replicates for validation. GC-MS analysis included six replicates.

## Acknowledgments

We acknowledge Dr. Kyle J. Roux for providing with Myc-BioID2 plasmid constructs, Dr. Swasti Raychaudhuri and the Proteomics facility, CCMB, Hyderabad, for peptide identification.

## Supplementary Materials

**Supplementary Fig. 1:**
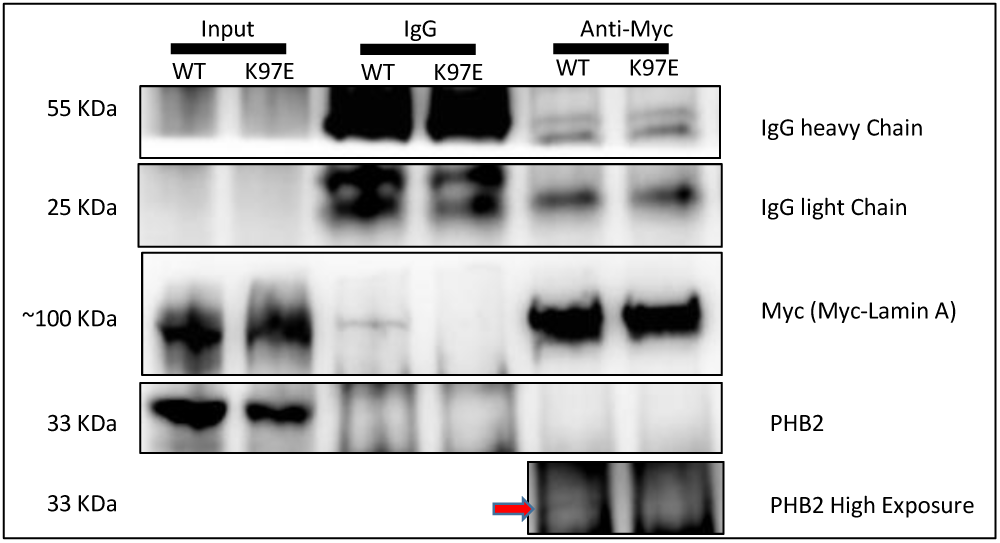
lamin A immunoprecipitation using myc-BioID2-lamin A constructs followed by immunoblotting using myc and PHB2 to confirm lamin A-PHB2 interaction.

**Supplementary Fig. 2:**
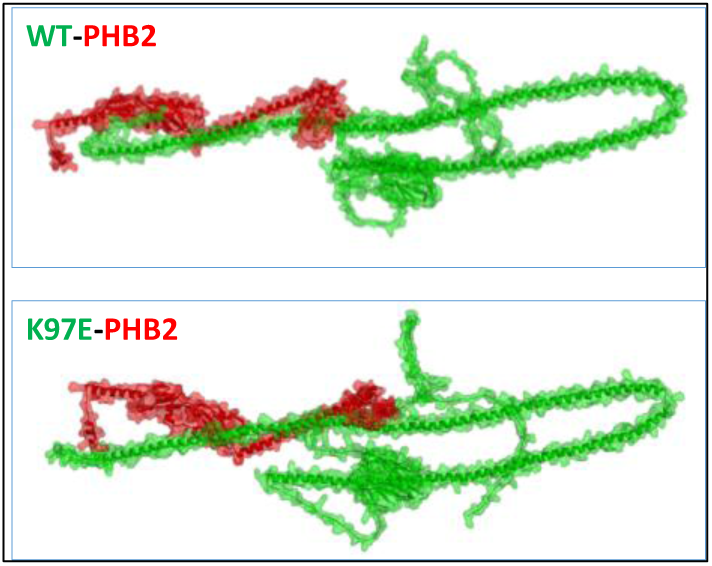
Molecular docking of PHB2 (red) and lamin A variants (green) showing interaction paradigm.

**Supplementary Fig. 3:**
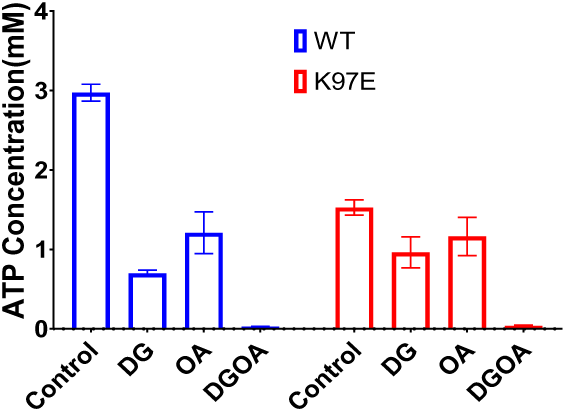
ATP concentration in control, 2-deoxyglucose (DG) treated, oligomycin (OA) treated, and DG and OA treated cells normalized with respect to MTT values.

**Supplementary Fig. 4:**
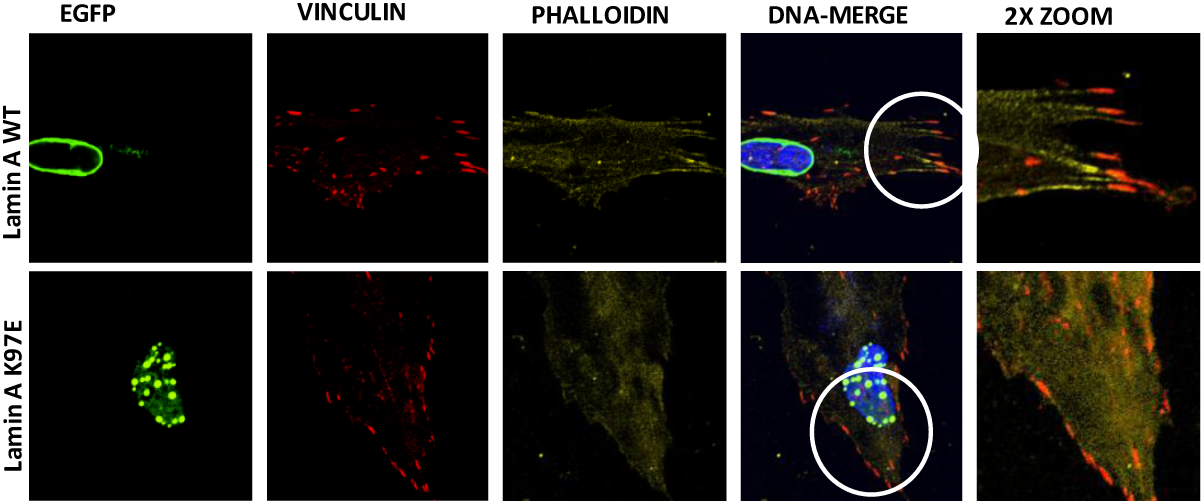
Immunofluorescence imaging showing vinculin morphology and distribution in transfected cells.

**Supplementary Fig. 5:**
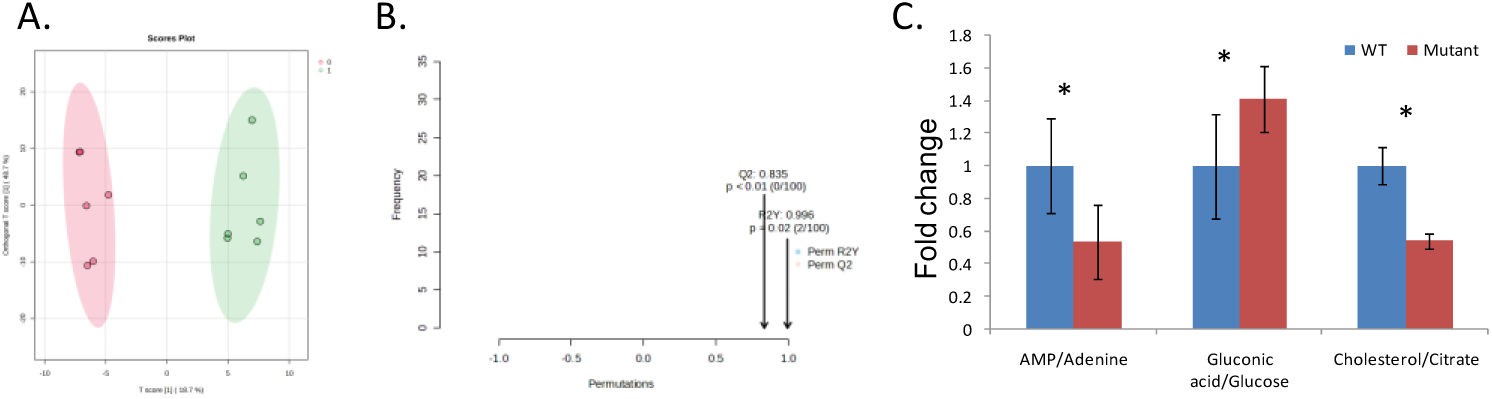
A. Supervised principal component analysis (PCA) of the K97E and WT metabolome. B. Permutation test. C. Quantified ratio of selected metabolites.

**Supplementary Fig. 6:**
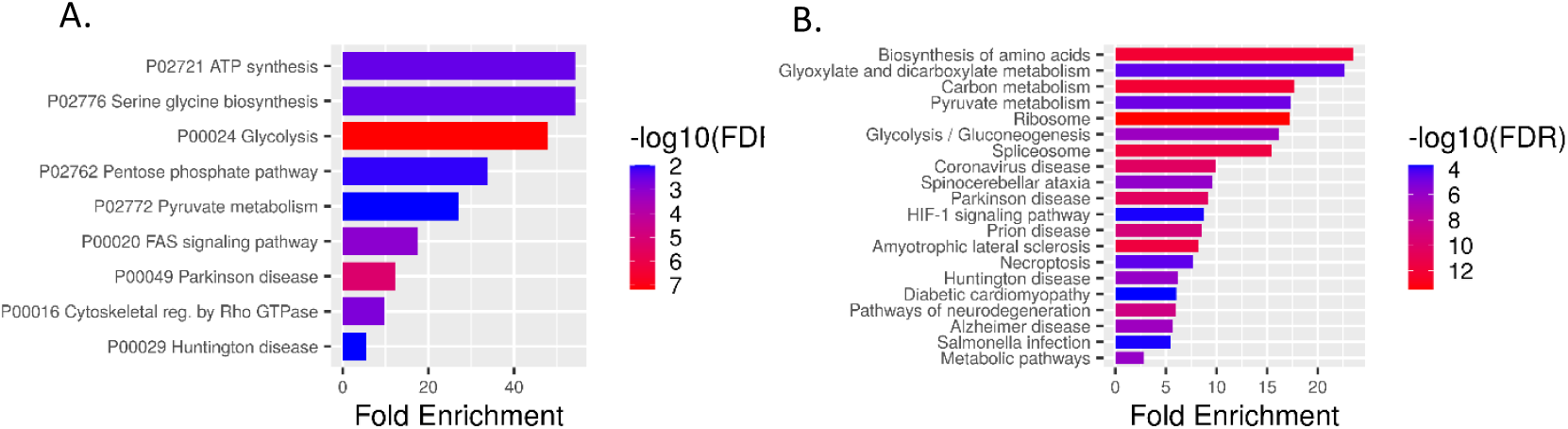
Enrichment of lamin A (WT) interactome using A. PANTHER and B. KEGG pathway database.

**Supplementary Fig. 7:**
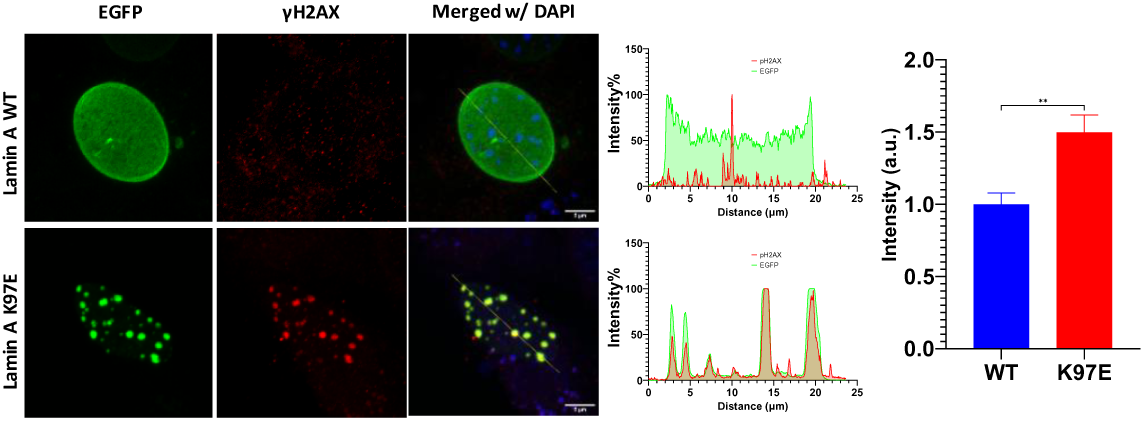
A. Immunofluorescence imaging showing pH2AX in nucleus of transfected cells. B. Plot profile showing pH2AX distribution. C. Quantitative analysis of pH2AX fluorescence intensity from confocal images.

**Supplementary table 1:**
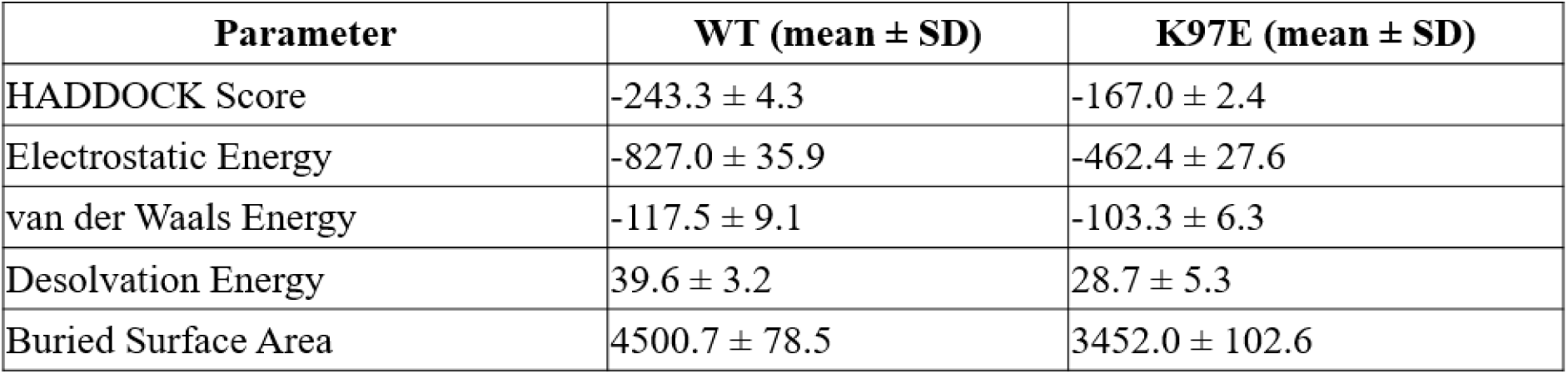
Energy parameters after docking and energy minimisation between PHB2 and lamin A WT/K97E.

| Parameter | WT (mean ± SD) | K97E (mean ± SD) |
| --- | --- | --- |
| HADDOCK Score | -243.3 ± 4.3 | -167.0 ± 2.4 |
| Electrostatic Energy | -827.0 ± 35.9 | -462.4 ± 27.6 |
| van der Waals Energy | -117.5 ± 9.1 | -103.3 ± 6.3 |
| Desolvation Energy | 39.6 ± 3.2 | 28.7 ± 5.3 |
| Buried Surface Area | 4500.7 ± 78.5 | 3452.0 ± 102.6 |

**Supplementary table 2:** Pathway analysis of altered metabolomic signature in K97E.

| Pathway | Match status | p-value | -Log(p) | FDR-adjusted |
| --- | --- | --- | --- | --- |
| Valine, leucine and isoleucine biosynthesis | 5/8 | 5.30E-07 | 6.2754 | 4.24E-05 |
| Alanine, aspartate and glutamate metabolism | 7/28 | 4.17E-06 | 5.3794 | 1.67E-04 |
| Galactose metabolism | 6/27 | 4.59E-05 | 4.3385 | 0.001223 |
| Citrate cycle (TCA cycle) | 5/20 | 1.16E-04 | 3.9354 | 0.002321 |
| Pantothenate and CoA biosynthesis | 4/20 | 0.0014939 | 2.8257 | 0.023903 |
| Arginine biosynthesis | 3/14 | 0.0050789 | 2.2942 | 0.067719 |
| Butanoate metabolism | 3/15 | 0.0062328 | 2.2053 | 0.071232 |
| Glyoxylate and dicarboxylate metabolism | 4/32 | 0.0088079 | 2.0551 | 0.088079 |
| Starch and sucrose metabolism | 3/18 | 0.010578 | 1.9756 | 0.094028 |
| Valine, leucine and isoleucine degradation | 4/14 | 0.019213 | 1.7164 | 0.15235 |
| Pyruvate metabolism | 4/23 | 0.020949 | 1.6788 | 0.15235 |
| One carbon pool by folate | 3/26 | 0.029116 | 1.5359 | 0.19411 |
| Glutathione metabolism | 3/28 | 0.035374 | 1.4513 | 0.21769 |

**Supplementary Methods Table 1:** List of Antibodies and fluorescent probes.

| Target | Catalogue | Company |
| --- | --- | --- |
| Anti Rabbit 546 | A11010 | Thermo Fischer Scientific |
| Anti-Mouse HRP | 32430 | Thermo Fischer Scientific |
| Anti-Mouse 546 | A11003 | Thermo Fischer Scientific |
| Anti-Mouse 633 | A21052 | Thermo Fischer Scientific |
| Anti-Rabbit 633 | A21070 | Thermo Fischer Scientific |
| Anti-Rabbit HRP | 32460 | Thermo Fischer Scientific |
| Arp2 | PA527879 | Thermo Fischer Scientific |
| Beta-Actin | A5316 | Sigma Aldrich |
| Drp1 | 8570 (D6C7) | Cell Signalling Technology |
| FAK | 396500 | Thermo Fischer Scientific |
| GAPDH | AB9485 | Abcam |
| INF2 | 20466-1-AP | proteintech |
| Lamin A/C | L1293 | Sigma Aldrich |
| Myc Tag | 71D10 | Cell Signalling Technology |
| Normal Rabbit IgG | 2729 | Cell Signalling Technology |
| Opa1 | 1E8 1D9 | Thermo Fischer Scientific |
| pERK1/2 | AB50011 | Abcam |
| Phalloidin-568 | A12380 | Thermo Fischer Scientific |
| PHB1 | PA519556 | Thermo Fischer Scientific |
| PHB2 | PA581532 | Thermo Fischer Scientific |
| p-Paxillin | 69363 (E9U9F) | Cell Signalling Technology |
| RhoA | 67B9 | Cell Signalling Technology |
| ROCK1 | PA522262 | Thermo Fischer Scientific |
| Vinculin | V9131 | Sigma Aldrich |

**Supplementary Methods Table 2:**
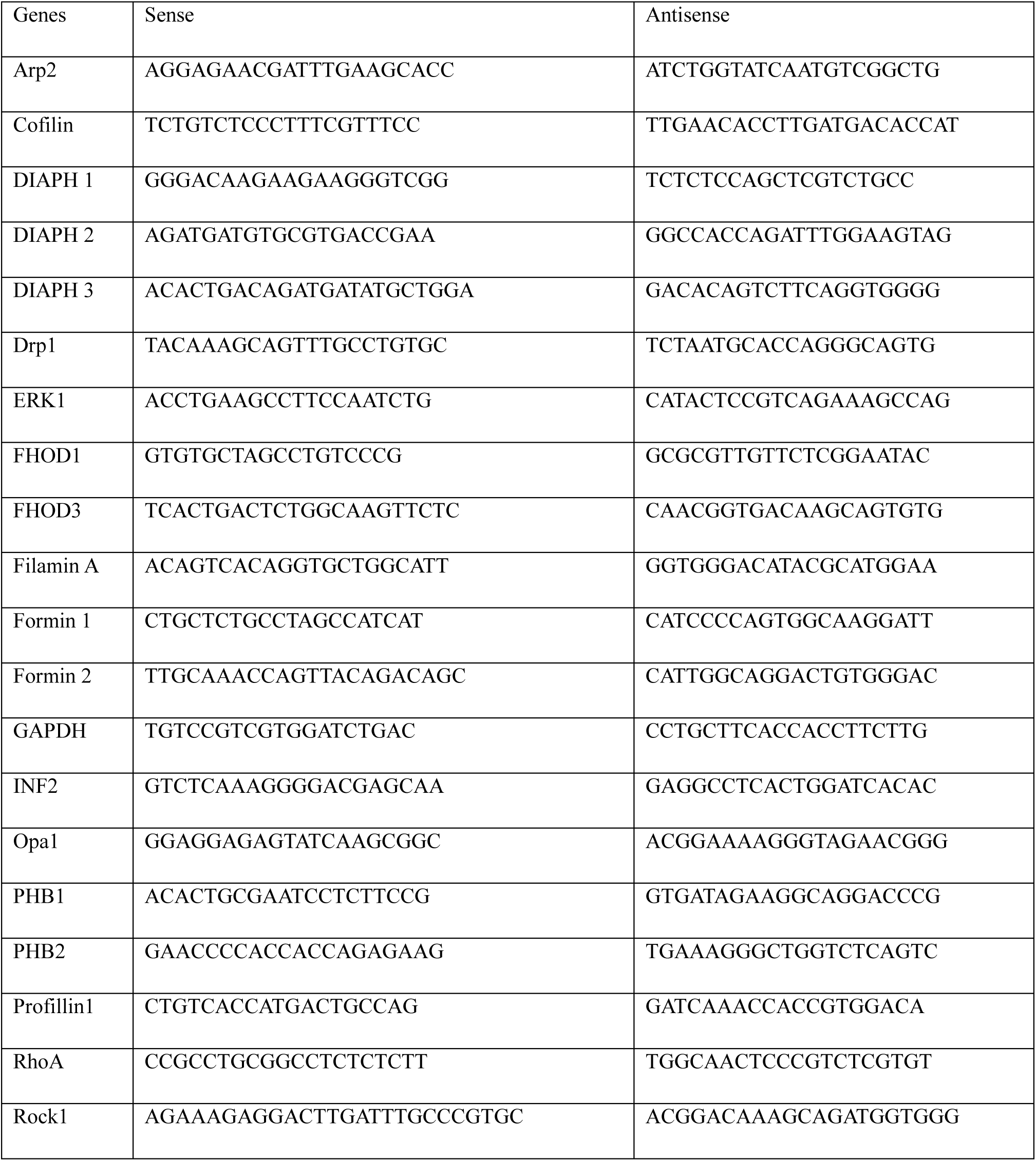
List of qPCR primers.

